# A new method to estimate the ecological niche through *n*-dimensional hypervolumes that combines convex hulls and elliptical envelopes

**DOI:** 10.1101/2022.03.03.482921

**Authors:** Jaime Carrasco, Fulgencio Lisón, Laura Jiménez, Andrés Weintraub

**Affiliations:** University of Chile, Industrial Engineering Department, Santiago, Chile; Complex Engineering System Institute - ISCI, Santiago, Chile; Wildlife Ecology and Conservation Lab, Departamento de Zoología, Facultad de Ciencias Naturales y Oceanográficas, Universidad de Concepción, Casilla 160-C, Concepción, Chile; Laboratorio de Ecología del Paisaje y Conservación, Departamento de Ciencias Forestales y Medioambiente, Facultad de Ciencias Agropecuarias y Forestales, Universidad de La Frontera, Box 54-D, Temuco, Chile; School of Life Sciences, University of Hawai’i at Mānoa, 2538 McCarthy Mall, Honolulu, HI 96822, USA

**Keywords:** bats, chiroptera, presence-only data, realized niche, niche overlap

## Abstract

1. Methods that estimate the niche of a species by calculating a convex hull or an elliptical envelope have become popular due to their simplicity and interpretation, given Hutchinson’s conception of the niche as an *n*-dimensional hypervolume.
2. It is well known that convex hulls are sensitive to outliers and do not have the ability to differentiate between regions of low and high concentration of presences, while the elliptical envelopes may contain large regions of niche space that are not relevant for the species. Thus, when the goal is to estimate the realized niche of the species, both methods may overestimate the niche.
3. We present a methodology that combines both the convex hull and the elliptical envelope methods producing an *n*-dimensional hypervolume that better fits the observed density of species presences, making it a better candidate to model the realized niche. Our method, called the CHE approach, allows defining regions of iso-suitability as a function of the significance levels inherited from the method (Mahalanobis distance model, minimum covariance determinant, or minimum volume ellipsoid) used to fit an initial elliptical envelope from which we then discard regions not relevant for the species by calculating a convex hull.
4. We applied the CHE approach to a case study of twenty-five species of bats present in the Iberian Peninsula, fitting a hypervolume for each species and comparing them to both the convex hulls and elliptical envelopes obtained with the same data and different values of *n*. We show that as the number of variables used to define the niche space increases, both the convex hull and elliptical envelope models produce overly large hypervolumes, while the size of the hypervolume fitted with the CHE approach remains stable. As a consequence, similarity measures that account for the niche overlap among different species may be inflated when using convex hulls or elliptical envelopes to model the niche; something that does not occur under the CHE approach.

## 1 Introduction

Since Hutchinson (1957) formalized the concept of the ecological niche as a *n*-dimensional hypervolume that contains the observations and conditions where a species adequately persists and survives, a variety of mathematical methods for its modeling have proliferated (see e.g. Blonder et al. (2014); Junker et al. (2016); Peterson et al. (2011)). These methods rely on a variety of approaches including traditional statistical techniques, computational geometry, convex analysis and machine learning algorithms (Blonder et al., 2018; Guo et al., 2005). At the same time, ecologists have been using the concept of hypervolume for a wide range of applications including niche modeling (Blonder et al., 2015; Cornwell et al., 2006; Soberón and Nakamura, 2009), to understand species coexistence and richness (Abrams, 1983; Rappoldt and Hogeweg, 1980), to study intra- and interspecific relationships (Pianka et al., 2017), to predict species invasions (Moles et al., 2008; Van Kleunen et al., 2010), in functional ecology (Lamanna et al., 2014; Swenson et al., 2012), to develop epidemiological and zoonotic models (Escobar, 2020) and to determine the state of stability at a community or ecosystem level (Barros et al., 2016). To define the niche of a species, the idea is to delineate a hypervolume as a geometrical shape within a multidimensional Euclidean space (or hyperspace) that can be used to describe the topology of the ecological system under study and its spatial relationship with other species (Blonder, 2018). In this context, extant methods to fit hypervolumes have being criticized because of their poor performance in high-dimensional spaces, or the high variability of their outputs according to the parameter choice (Junker et al., 2016; Etherington, 2021). Thus, there is a need to explore and compare different methods to estimate *n*-dimensional hypervolumes used to model the realized niche of a species.

In the fields of species distribution modeling (SDM) and ecological niche modeling (ENM), there is a variety of methods for delineating a hypervolume ranging from fitting polygons (Busby, 1991; Broennimann et al., 2007; Cornwell et al., 2006; Gruson, 2020) to more complex techniques based on Kernel density estimates (Blonder et al., 2014; Broennimann et al., 2012; Blonder et al., 2018). However, in each of these fields, the fitted hypervolumes are used to understand and analyze different objects (Peterson and Soberón, 2012; Warren, 2012). In SDM, the shape of the hypervolume is usually not relevant since the aim is to delimit the current geographic range of the species. As a result, some modeling approaches estimate objects in an *n*-dimensional space which can be unbounded volumes (e.g., random forest, generalized linear models, boosted regression trees), for which an additional criterion needs to be applied to decide where the species’ distribution ends (binarization). On the other hand, several authors suggest that species’ fundamental climatic niches are convex in shape (Peterson et al., 2011; Soberón and Nakamura, 2009; Soberón and Peterson, 2020), and thus, in the field of ENM, methods that construct convex hypervolumes, or envelopes, have been preferred and widely applied (Pearce and Boyce, 2006).

The intuition of defining limits for each of the environmental variables relevant for the survival of the species captures the sense of a niche since the occurrence of species should be limited by these environmental factors. Therefore, the identification of the fundamental niche of a species, or any subset of it, can be achieved by building a convex set enclosing the occurrences of the species or a portion of them. However, if the goal is to reconstruct the fundamental niche of the species, using occurrence data is not enough (Jiménez et al., 2019). The sites where the species is observed as present come from the realized niche of the species, this is, they are determined not only by abiotic factors but are also influenced by biotic interactions and the dispersal limitations of the species. Thus, unless some supplementary information is contained in the data, we should not expect models to identify the whole breadth of the fundamental niche (Qiao et al., 2018). Here, we focus on methods to estimate convex hypervolumes from presence-only data as models for the realized niche of the species.

From the long list of methods to estimate convex hypervolumes used in SDM and ENM, the following stand out: (*1*) convex hulls or polyhedra surrounding all the occurrences (Broennimann et al., 2007; Cornwell et al., 2006; Dallas et al., 2017; Walker and Cocks, 1991); (*2*) elliptical envelopes which may contain all or a portion of the occurrences (Swanson et al., 2015; Etherington, 2021), and (*3*) hyper-rectangles or box-shaped approaches (Busby, 1991; Nix et al., 1986). It has been shown that there is no clear “best” way to delineate hypervolumes (Blonder, 2018; Merow et al., 2014) and the choice of an appropriate method will depend on the goals of the analysis and data limitations (Blonder et al., 2018; Etherington, 2021; Peterson et al., 2011). The convex hull (CH) approach uses all observations to build the minimum convex polyhedron that contains them. An unfavorable feature of the CH is that its topology is strongly influenced by peripheral observations, making it very sensitive to outliers. Furthermore, the CH approach does not identify suitability levels that could occur internally (Cerdeira et al., 2018), the fitted hypervolume is fixed and unique, so it does not distinguish among different confidence levels (Blonder et al., 2014). However, this approach has the advantage of being distribution-free, or non-parametric, because no assumption about the statistical distribution of the data is assumed. In contrast, the hypervolumes estimated through elliptical envelope (EE) methods are often obtained at different confidence levels, or incorporate different proportions of the observed occurrences excluding the farthest outliers. The EE approaches are parametric and they assume a symmetrical response to environmental limits, although there are situations in which this geometry does not fit the data well and therefore does not adequately represents a species niche (Austin, 1987). Additionally, the EE approaches are closely related to the Mahalanobis distance model which delineates ellipsoids that represent the geometric shape of the data points that are at a certain distance from an *n*-dimensional vector and incorporates the correlation between environmental variables through the estimation of a matrix that describes the orientation and extent of the niche (Farber and Kadmon, 2003; Clark et al., 1993; Etherington, 2021). Under this approach, the calculation of intersections between *n*-dimensional ellipsoids (which is often desired in community ecology), require the application of approximation methods that involve great computational efforts (Qiao et al., 2016; Rabiei and Saleeby, 2021; Swanson et al., 2015). Finally, the box-shaped approaches rely on the assumption that environmental variables act independently on the species, an assumption that is considered unrealistic (Maguire, 1973), and it is leading researchers to adopt more complex modelling approaches. For this reason, we focus on the first two approaches.

How these hypervolumes are built and how we can use them describe of the species’ niche is an important issue for fields such as macroecology and biogeography. From the mathematical and conceptual formalism presented by Blonder (2018), hypervolumes are defined as bounded geometric shapes. Following this concept, we developed an approach that combines the Convex Hull and the traditional Elliptical envelope methods. This method produces polyhedral hypervolumes based on presence-only data and it allows to define regions of iso-suitability whose intersection or overlap regions can be determined in an efficient and more representative manner. We hypothesized that the combination of CH and EE methods can fit a *n*-dimensional hypervolume that is more adjusted to the observations, better representing the geometric shape of the ecological niche, and remains aligned with Hutchinson’s niche concept. This work aims to: i) model the realized niche of a species by combining the convex and elliptical envelope methods; ii) compare the performance between the hypervolumes fitted with our proposed methodology and the ones fitted with the convex hull and elliptical envelop methods; and iii) discuss the advantages of our methodology and its applications in estimating niche overlap and species distributions.

## 2 Materials and Methods

### 2.1 Presence data and environmental variables

We used presence records of twenty-five bat species present in the Iberian peninsula (the species scientific names and number of observations can be found in Table 2). The Iberian peninsula has an approximate area of six thousand square kilometers, which we divided into 10 × 10 *km*^2^ UTM cells. These presence samples were obtained from the distribution maps “Atlas y Libro Rojo de los Mamíferos Terrestres de España” (Palomo et al., 2007) and “Atlas dos morcegos de Portugal” (Rainho et al., 2013), both recently updated with new records at the same resolution from personal surveys and references (Lisón et al., 2015; Lisón and Sánchez-Fernández, 2017). We only used those bat species with more than thirty records, and we did not include the twin species *Myotis nattereri* and *Myotis escalerai* because there is a lot of uncertainty regarding their distributions (Palomo et al., 2007; Juste et al., 2018).

We included two types of environmental variables in our analyses: climatic and land-use variables. Land-use variables were obtained following the procedure described in Lisón et al. (2015) and Lisón and Sánchez-Fernández (2017). We employed the land-use map provided by the CORINE Land Cover Project (see www.eea.europa.eu). Also, we constructed habitat-suitability models for bat species using as environmental variables the land-use cover data from the CORINE Project for 2006 (https://land.copernicus.eu/pan-european/corine-land-cover/clc-2006). Land-use cover was reclassified from the 44 initial categories to 16 definitive ones (see Lisón and Sánchez-Fernández (2017)). For each such category, we calculated the percentage of surface area included in each cell. We used six categories of land-use cover (forest, shrubland, pasture, wetland, irrigate and urban) because they are the most significative for bat in several studies (Lisón and Calvo, 2013; Smeraldo et al., 2018). In addition, we extracted four environmental variables from BIOMOD for use with each cell following Barbosa et al. (2012) (Table 1).

**Table 1:** Description of environmental variables used in our models for all the bat species.

| Drivers | Variable name | Description |
| --- | --- | --- |
| Climate | tmed | Average annual temperature (°C) |
|  | tjan | Average temperature in January (°C) |
|  | tjul | Average temperature in July (°C) |
|  | prec | Mean annual precipitation (mm) |
| Land cover | Forest | Forest land [%] |
|  | Shrubland | Shrubland [%] |
|  | Pasture | Pasture land [%] |
|  | Wetland | Wetland [%] |
|  | Irrigatedcrops | Irrigate crops [%] |
|  | Urban | Urban land [%] |

**Table 2:** Volumes of the fitted hypervolumes of each species of bats, with *p* = 0.95 and for each estimation approach.

| Species | Index | $R_\beta$ | Dim. | Hypervolumes | | | | | | | |
| --- | --- | --- | --- | --- | --- | --- | --- | --- | --- | --- | --- |
|  |  |  |  | MVEE | CH | COV | MCD | MVE | CH-COV | CH-MCD | CH-MVE |
| *Barbastella barbastellus* | 1 | 349 | $n = 3$<br>$n = 4$ | 263.69<br>1756.1 | 104.45<br>292.63 | 132.8<br>592.24 | 119.03<br>119.03 | 120.81<br>360.14 | 58.79<br>110.55 | 54.06<br>64.87 | 54.06<br>77.87 |
| *Eptesicus isabellinus* | 2 | 139 | $n = 3$<br>$n = 4$ | 230.42<br>2115.4 | 67.87<br>186.71 | 109.2<br>513.09 | 70.71<br>70.71 | 72.05<br>159.22 | 30.99<br>44.87 | 22.29<br>20.96 | 22.79<br>19.36 |
| *Eptesicus serotinus* | 3 | 776 | $n = 3$<br>$n = 4$ | 396.39<br>4369.7 | 158.88<br>656.74 | 155.84<br>839.42 | 132.92<br>132.92 | 133.81<br>516.93 | 73.53<br>175.45 | 61.54<br>123.5 | 67.44<br>140.87 |
| *Hypsugo savii* | 4 | 460 | $n = 3$<br>$n = 4$ | 217.6<br>1743.6 | 97.41<br>341.85 | 111.33<br>492.18 | 104.74<br>104.94 | 104.01<br>323.51 | 51.57<br>99.5 | 51.58<br>72.02 | 51.78<br>78.99 |
| *Myotis bechsteinii* | 5 | 278 | $n = 3$<br>$n = 4$ | 97.19<br>450.32 | 39.48<br>74.43 | 87.22<br>291.47 | 85.25<br>85.25 | 83.03<br>206.71 | 26.56<br>38.82 | 27.32<br>24.34 | 27.32<br>29.35 |
| *Myotis blythii* | 6 | 278 | $n = 3$<br>$n = 4$ | 288.42<br>2283.2 | 96.37<br>398.98 | 130.85<br>734.04 | 120.48<br>120.48 | 119.52<br>343.81 | 54.98<br>102.07 | 52.61<br>53.23 | 53.19<br>60.79 |
| *Myotis capaccinii* | 7 | 94 | $n = 3$<br>$n = 4$ | 81.23<br>168.48 | 24.37<br>20.15 | 63.01<br>116.97 | 53.85<br>53.85 | 52.61<br>93.1 | 15.69<br>11.93 | 13.33<br>9.4 | 13.33<br>8.68 |
| *Myotis daubentonii* | 8 | 859 | $n = 3$<br>$n = 4$ | 279.1<br>2654.8 | 141.09<br>555.73 | 150.38<br>708.49 | 140.08<br>140.08 | 140.93<br>478.3 | 78.94<br>181.38 | 72.82<br>118.9 | 74.67<br>141.35 |
| *Myotis emarginatus* | 9 | 363 | $n = 3$<br>$n = 4$ | 129.85<br>581.71 | 64.03<br>161.89 | 113.97<br>448.04 | 114.34<br>115.28 | 113.62<br>346.08 | 56.17<br>96.75 | 55.91<br>65.81 | 51.79<br>84.19 |
| *Myotis myotis* | 10 | 811 | $n = 3$<br>$n = 4$ | 314.25<br>2683.2 | 131.7<br>604.38 | 139.23<br>730.47 | 127.23<br>127.2 | 130.89<br>438.59 | 70.37<br>179.38 | 63.1<br>100.51 | 62.98<br>110.84 |
| *Myotis mystacinus* | 11 | 101 | $n = 3$<br>$n = 4$ | 120.81<br>503.16 | 42.15<br>84.37 | 88.31<br>366.96 | 79.24<br>79.24 | 79.96<br>271.26 | 28.48<br>43.37 | 25.49<br>39.06 | 24.55<br>40.26 |
| *Miniopterus schreibersii* | 12 | 810 | $n = 3$<br>$n = 4$ | 482.65<br>5726.3 | 169.9<br>795.29 | 140.62<br>757.11 | 130.11<br>130.16 | 130.89<br>449.38 | 70.9<br>183.67 | 68.66<br>97.53 | 69.64<br>108.12 |
| *Nyctalus lasiopterus* | 13 | 208 | $n = 3$<br>$n = 4$ | 455.88<br>3810.2 | 155.25<br>571.96 | 219.08<br>1271.1 | 169.28<br>169.28 | 170.58<br>447.84 | 81.27<br>150.06 | 59.61<br>53.01 | 56.66<br>67.77 |
| *Nyctalus leisleri* | 14 | 620 | $n = 3$<br>$n = 4$ | 533.44<br>5377.3 | 164.54<br>748.45 | 147.78<br>741.98 | 126.35<br>126.35 | 128.77<br>443.16 | 81.1<br>193.15 | 66.88<br>103.11 | 67.85<br>124.66 |
| *Plecotus austriacus* | 15 | 294 | $n = 3$<br>$n = 4$ | 177.46<br>1193.7 | 82.78<br>265.48 | 102.35<br>534.27 | 92.15<br>92.04 | 91.64<br>395.28 | 43.2<br>94.73 | 38.7<br>65.82 | 43.09<br>70.44 |
| *Plecotus auritus* | 16 | 741 | $n = 3$<br>$n = 4$ | 340.31<br>2861.1 | 145.27<br>613.71 | 125.38<br>625.54 | 114.18<br>114.21 | 116.33<br>342.14 | 67.46<br>157.59 | 62.64<br>82.04 | 59.52<br>94.26 |
| *Pipistrellus kuhlii* | 17 | 1095 | $n = 3$<br>$n = 4$ | 661.71<br>8491 | 225.29<br>1027.8 | 157.16<br>807.38 | 130.21<br>130.08 | 130.72<br>438.67 | 78.94<br>208.45 | 69.77<br>83.31 | 74.33<br>129.68 |
| *Pipistrellus nathusii* | 18 | 33 | $n = 3$<br>$n = 4$ | 89.96<br>348.74 | 27.39<br>53.67 | 127.49<br>802.38 | 126.56<br>126.56 | 110.52<br>361.54 | 23.95<br>28.74 | 16.42<br>12.06 | 14.29<br>17.67 |
| *Pipistrellus pipistrellus* | 19 | 2297 | $n = 3$<br>$n = 4$ | 752.29<br>10010 | 275.01<br>1501.1 | 173.83<br>917.85 | 148.33<br>148.33 | 151.13<br>558.78 | 98.4<br>294.35 | 80.94<br>152.14 | 85.76<br>183.74 |
| *Pipistrellus pygmaeus* | 20 | 1938 | $n = 3$<br>$n = 4$ | 692.91<br>9453.7 | 264.91<br>1325.3 | 167.3<br>887.24 | 144.73<br>144.28 | 144.69<br>538.85 | 90.37<br>256.86 | 77.14<br>133.91 | 78.99<br>165.88 |
| *Rhinolophus euryale* | 21 | 522 | $n = 3$<br>$n = 4$ | 256.4<br>2102.6 | 111.07<br>432.25 | 139.55<br>672.21 | 134.19<br>134.19 | 135.01<br>465.05 | 74.14<br>168.63 | 65.55<br>98.63 | 65.22<br>107.83 |
| *Rhinolophus ferrumequinum* | 22 | 1531 | $n = 3$<br>$n = 4$ | 421.91<br>3036.7 | 169.98<br>870.32 | 156.13<br>777.51 | 145.82<br>139.5 | 147.34<br>503.8 | 90.97<br>231.32 | 80.62<br>141.37 | 82.24<br>168.76 |
| *Rhinolophus hipposideros* | 23 | 1298 | $n = 3$<br>$n = 4$ | 227.73<br>1805.1 | 82.67<br>723.98 | 95.96<br>720.3 | 66.57<br>145.8 | 70.38<br>473.47 | 35.2<br>220.4 | 26.94<br>134.21 | 29.16<br>157.62 |
| *Rhinolophus mehelyi* | 24 | 214 | $n = 3$<br>$n = 4$ | 227.73<br>1805.1 | 82.67<br>280.39 | 95.96<br>486.54 | 66.57<br>66.57 | 70.38<br>235 | 35.2<br>67.7 | 26.94<br>30.38 | 29.16<br>33.88 |
| *Tadarida teniotis* | 25 | 1080 | $n = 3$<br>$n = 4$ | 564.57<br>7081.8 | 207.11<br>1044 | 156.43<br>857.29 | 134.36<br>134.38 | 136.27<br>529.82 | 82.41<br>223.83 | 70.51<br>115.53 | 73.93<br>155.47 |
| | | AVG : | $n = 3$<br>$n = 4$ | 339.29<br>3388.4 | 128.97<br>545.26 | 133.7<br>667.68 | 118.01<br>118.03 | 118.22<br>388.82 | 62.04<br>142.54 | 54.625<br>79.826 | 55.43<br>95.133 |
| | | STD : | $n = 3$<br>$n = 4$ | 193.69<br>2846.4 | 69.514<br>396.93 | 33.053<br>233.87 | 28.661<br>28.638 | 29.032<br>122.42 | 24.125<br>77.679 | 21.321<br>42.607 | 22.124<br>52.526 |

Instead of working with the raw environmental variables mentioned above, we applied a principal component analysis to reduce the dimensionality of the multivariate space where the hypervolumes are defined, and to work with orthogonal variables. In Fig. S1, we show a scree plot of the percent variability explained by each principal component. This plot only shows the first seven principal components, which explain about 90% of the total variance. The first component by itself explained less than 40% of the variance, so more components need to be considered. The first three components explain 55.17% of all variability, and the first four over 65%. We worked with multivariate environmental spaces from two to five dimensions using the first five principal components.

### 2.2 Methods for delimiting hypervolumes

Here, we first outline different methods to estimate a hypervolume in an *n*-dimensional space from a set of presence data, and we explain how they are used to delimit the niche of the species. Then, we introduce a new method for delimiting hypervolumes which combines two existing approaches. Finally, we applied all these methods to the assembled dataset of bat species and compared their outputs. However, before continuing, we introduce the basic notation and terminology which will be used throughout of the next sections.

Let 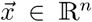 be a vector whose entries represent the environmental conditions observed at a site where a species of interest is observed as present and which are described in Table 1. As we mentioned in the previous section, we will specifically use the transformed variables using PCA and not the original variables to achieve orthogonality (then *n* ≤ 7). Let ℐ be the cindex set of all the cells in the study area and 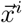 be vector of variables measured in cell *i*. In our dataset, the study area contains 6,169 cells (ℐ = {1, …, 6, 169}), in which the twenty-five species of bats are distributed. We denote by ℬ = {1, …, 25} the index set of the bat species so that *β* ∈ ℬ represents a particular bat species (see the second column in Table 2), and *R*_*β*_ represents the number of records/presences of *β* in the study area. Finally, ℐ_*β*_ is the subset of cell indices where the specie *β* is present. Thus, 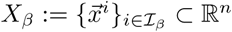 denotes the sample of observations of species *β*.

#### 2.2.1 Convex hulls (CH-approach)

The estimation of an *n*-dimensional hypervolume from a set of observations *X*_*β*_ can be achieved by fitting a convex hull (CH), defined as the smallest convex set containing *X*_*β*_ and denoted by *CH*(*X*_*β*_). If the set of points (observations) is finite, it can be proved that convex hull is a polytope, or a bounded polyhedron. By definition, a polyhedron is a set that can be written as a linear system of inequalities: 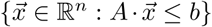, where *A* is a *m* × *n* matrix, *b* is a vector in ℝ^*m*^ for certain positive integer *m*, and 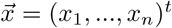 is an *n*-dimensional vector. Thus, *CH*(*X*_*β*_) depends on two parameters: a matrix *A* and a vector *b*, which are fully determined by the sample of presences.

The polyhedral representation of the CH is important because it helps calculating its volume by applying the classical *n*-simplex formula (see for example Allgower and Schmidt (1986)). In our study, we computed the CH and its volume using the QuickHull C++ interface for MATLAB (Barber et al., 1996).

#### 2.2.2 Elliptical envelopes

Formally, a multidimensional ellipsoid ℰ can be written as follows:

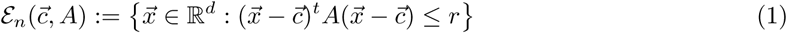

where *A* is a symmetric positive definite matrix (*A ≻* 0) of order *n* × *n* and 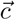 represents its geometric center. It is important to note that from the ellipsoid formulation (Eq. (1)), *r* is a constant and, without loss of generality, we could fix *r* = 1 by rescaling the matrix *A*.

Before describing some extent approaches to fit ellipsoids, or elliptical envelope (EE) methods, let us introduce the formula to calculate the volume of an ellipsoid, which will be used to compare the elliptical envelopes estimated with the different approaches:

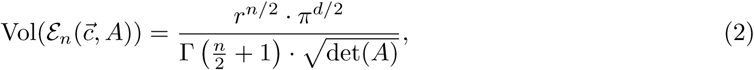

where det(·) denotes the matrix determinant and Γ(·) is the gamma function (see Wilson (2010)).

There exist different methods to determine an ellipsoid that enclose all or a portion of the observations in a sample, below we briefly describe the most popular ones.

##### Minimum-Volume Enclosing Ellipsoid

The minimum-volume enclosing ellipsoid (MVEE) is defined as the smallest ellipsoid (of minimum volume) that contains all the observations. Formally, from Eqs. (1) and (2) the problem of finding the MVEE of the set 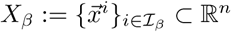 can be formulated as:

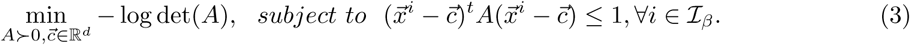

We solved the optimization problem of the Eq. (3) using the SDPT3 MATLAB-package (Tütüncü et al., 2003).

##### Sample mean and covariance: the Mahalanobis distance model

Let 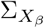 and 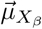 be the sample covariance matrix and the sample mean vector of the dataset *X*_*β*_, respectively. If we set 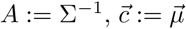 in Eq. (1), then, for a given value of 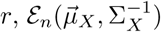 defines an ellipsoid around the average environmental conditions observed in the sample. In previous works, this method to estimate the parameters of an ellipsoid has being called the Mahalanobis distance model (Farber and Kadmon, 2003). Here, we refer to this method as the COV-approach.

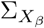 and 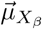 can be interpreted as the maximum likelihood parameter estimates of a multivariate normal distribution. In other words, if we assume that the random sample *X*_*β*_ comes from a multivariate normal random model, the values of the parameters that maximize the probability of observing that particular sample are the sample mean and the sample variance-covariance matrix. The assumption that the sample comes from a normal distribution may be strong, however, it allow us to determine a value of *r* that defines an ellipsoid as a confidence region. Under this assumption, the quadratic form 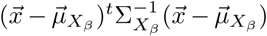 follows a chi-squared distribution with *n* degrees of freedom. Let 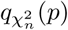 be the quantile function of the chi-square distribution with *n* degrees of freedom where *p* can be any value in the interval (0, 1), by setting 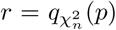 in Eq. (1), we get the ellipsoid 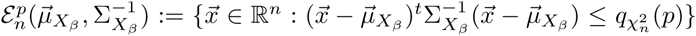 which corresponds to the confidence region of probability 100(1 − *α*)%, where *α* is a significance level. For example, the significance level *α* = 0.05 corresponds to a 95% confidence level such that the ellipsoid 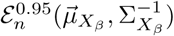 will contain approximately 95% of the presence points of *X*_*β*_, under the hypothesis that the sample comes from a multivariate normal distribution with parameters 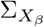 and 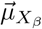.

##### Minimum volume ellipsoid and minimum covariance determinant

Robust estimates of the center 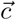 and the scatter matrix *A* of the sample *X*_*β*_ can be obtained by the minimum covariance determinant (MCD) estimator (Rousseeuw and Driessen, 1999) or minimum-volume ellipsoid (MVE) (Rousseeuw, 1985). Both methods use a number *h* ≤ *R*_*β*_ observations, where the value of *h* can be chosen by the user and it determines the robustness of the resulting estimator, however, they differ in how the estimates of the matrix A are calculated. The MCD estimates the matrix with smallest covariance determinant, while the MVE constructs an ellipsoid that encloses the *h* observations, whose matrix shape is of minimum volume. Therefore, both methods can be seen as special case of the COV and MVEE approaches, respectively.

The two methods are very useful for outlier detection in multivariate datasets (Van Aelst and Rousseeuw, 2009) and have been used in different applications (see e.g. Escobar et al. (2018), Qiao et al. (2016), Yañez-Arenas et al. (2018), or Torrejón-Magallanes et al. (2021). We used the FAST-MCD and FAST-MVE algorithms implemented in the FSDA library (Riani et al., 2012) to compute both estimators, obtaining the ellipsoids 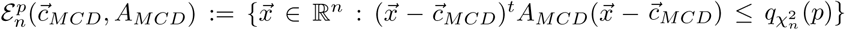 and 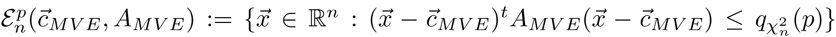, respectively. In both cases, we set *h* = [*R*_*β*_ + *d* + 1] because this produces more robust estimators (Hubert and Debruyne, 2010; Van Aelst and Rousseeuw, 2009).

#### 2.2.3 Combining the convex hulls and envelope ellipsoids: a new model for the realized niche

Recent studies have suggested that modeling approaches based on elliptical envelopes may be more appropriate when the goal is to estimate the fundamental niche of a species (Soberón and Peterson, 2020; Etherington, 2021); while the realized niche may be modeled through more complex shapes (Jiménez-Valverde et al., 2008). On the other hand, it is well known that the CH approach is sensitive to outliers, which may be common in presence-only samples even after going through standard data cleaning procedures. Therefore, none of these methods by themselves seem to provide a satisfactory representation of the realized niche of a species.

Still, we should recognize that these approaches posses particular strengths including that both elliptical envelopes and convex hulls are easily estimated from a sample of observations *X*_*β*_, using the functions provided in different scientific computing software. We thus propose to combine both approaches into a single improved method that aims to delimit the realized niche of a species as a hypervolume in an *n*-dimensional space as follows. First, we estimate an elliptical envelope 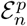 with the sample *X*_*β*_ applying any of the methods described in the Section 2.2.2, except the MVEE because it does not depend on *p* (it contains all the observations). Second, we use the subset of observations that fall inside the ellipsoid 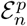, denoted by 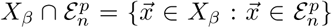, to build a CH which we denote by

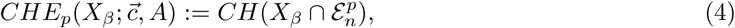

where 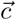 and *A* are the estimated center and shape matrix, respectively, and the acronym CHE stands for Convex Hull and Ellipsoid, as the new method combines both approaches. Depending on the elliptical envelope method used in the first step of our method (COV, MCD or MVE), the parameters are substituted by 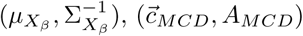, or 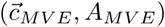, accordingly.

### 2.3 Comparative analysis

In order to compare the different approaches to delineating the realized niche, we proceeded as follows. First, we fit models of the niche of each bat species with the different approaches presented above, for the *n* = 3 and *n* = 4, and using a confidence level of 95% (or *p* = 0.95); and second, we perform a sensitivity analysis of the size of the hypervolumes as a function of the variation of the hyperspace dimension and the significance value.

## 3 Results

### 3.1 Comparison among different methods to estimate hypervolumes

To illustrate the differences between the four EEs approaches described in Section 2, we used the data corresponding to species *Barbastella barbastellus*, which we refer to as Bbar and whose corresponding sample of presences is *X*_1_ (*R*_*β*_ = 349). In Fig. 1, the enclosing ellipsoid generated by the MVEE, COV, MCD, and MVE methods are presented for the dimensions *n* = 2 (left chart) and *n* = 3 (right chart), which where calculated for *p* = 0.95. In both charts, we can observe that MVEE is an ellipsoid that encloses all the data points, while the remaining ones leave points outside the hypervolume. Also, in the 2D representation, no significant differences are seen between the hypervolumes built by COV, MCD and MVE. In the 3D case, COV generates more spherical shapes, unlike MCD and MVE whose corresponding ellipsoids are more elongated, and enclose the region with the highest concentration of observations.

**Figure 1:**
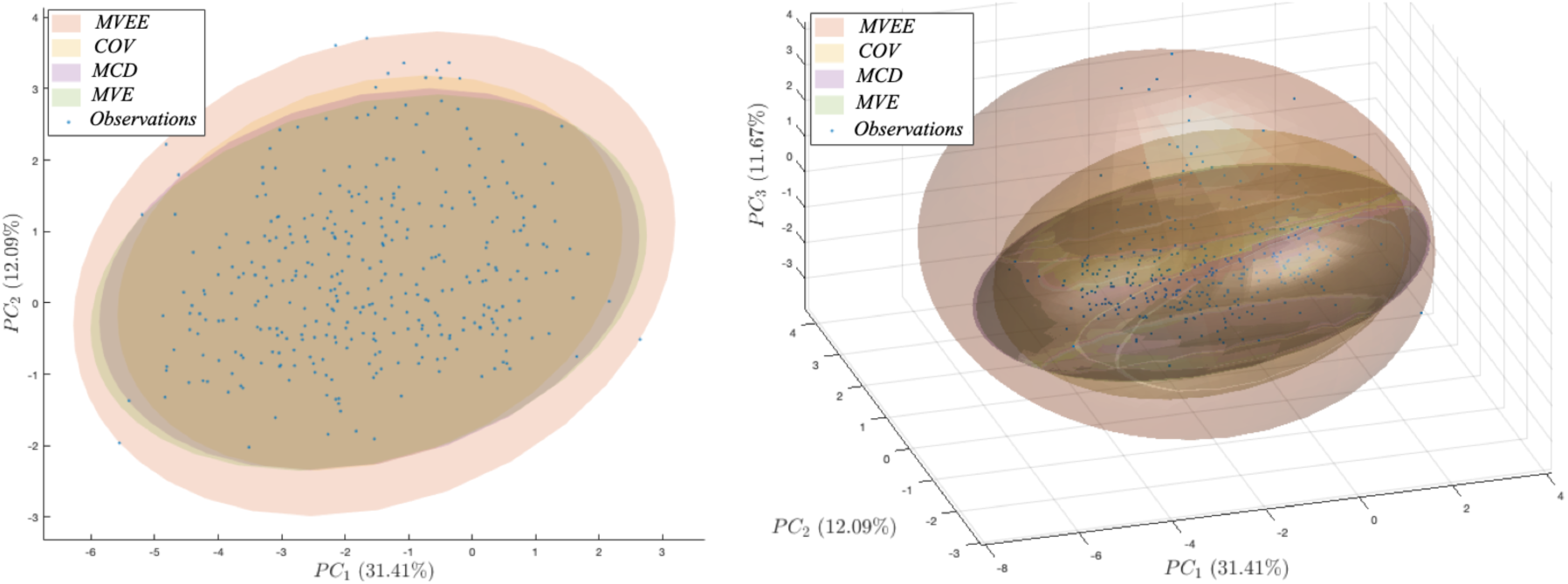
Enclosing ellipsoid generated by the MVEE, COV, MCD and MVE approaches using the Bbar species observations (blue dots), with *p* = 0.95, *n* = 2 (left chart) and *n* = 3 (right chart).

Similarly, in the 3D case (left panel in Fig. 2), we show the hypervolumes obtained from the MVEE and CH methods. We observe that the hypervolume estimated through CH is contained within the ellipsoid estimated through MVEE, because, by definition, the CH is the minimum convex set that contains the data set *X*_1_. As explained above, the *CH*(*X*_1_) can be expressed using a set of linear inequalities, and these are implicitly represented in the facets of the figure. In the middle panel of Fig. 2, we show the hypervolumes obtained with the methods CH and CH-COV are drawn for *p* = 0.95 (i.e., *CHE*_.95_(*X*_1_)). We can see that the CH polyhedron contains all the observations and is larger than the polyhedron obtained with the CH-COV method. Note also that the CH-COV method eliminates outliers using the Mahalanobis distance, and then builds the convex hull with the remaining points, producing a more limited hypervolume which is less sensitive to outliers and errors. Finally, the right chart in Fig. 2 shows the contraction of the polyhedrons estimated by the CH-COV method for *p* = 0.95 and CH-COV for *p* = 0.9, which highlights the ability of the CHE approach to build hypervolumes that better enclose the highest concentration of observed presences, a desirable features when modeling the realized niche of a species.

**Figure 2:**
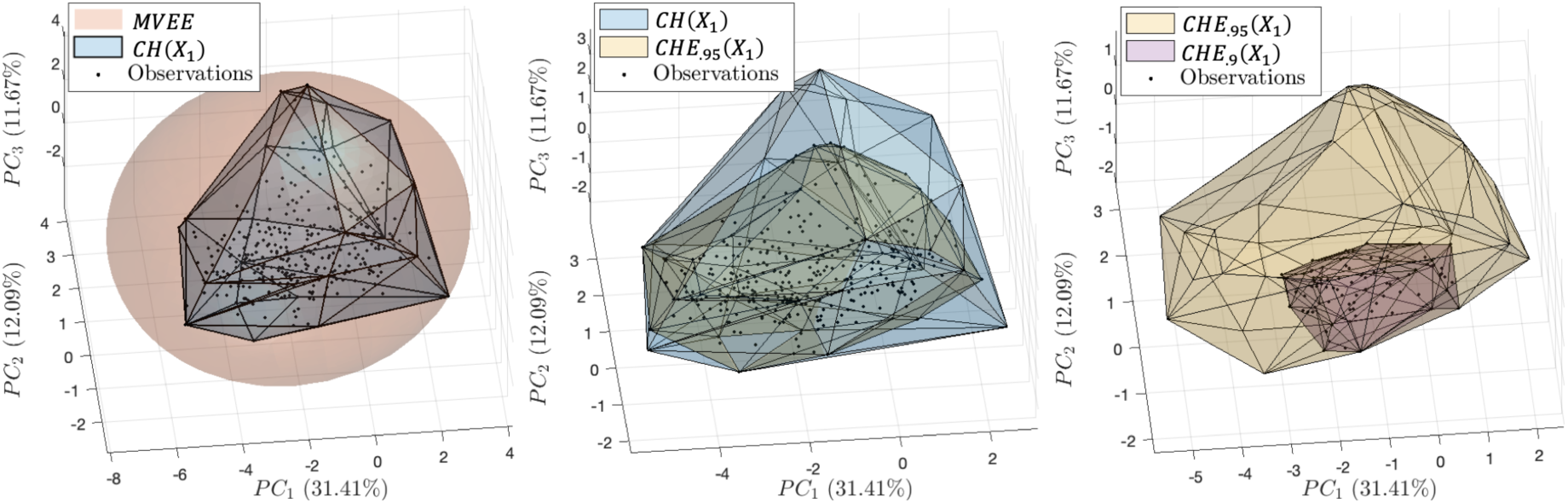
Comparison of the hypervolumes generated by the MVEE, CH, and CHE approaches for the species Bbar with *n* = 3.

In Table 2, we report the volume values obtained for each method for *n* = 3 and *n* = 4. respectively. In Fig. 3 we present the associated boxplots for a graphical understanding. The first four columns in this table, we show the volume of the hypervolumens obtained with the MVEE, COV, MCD and MVE methods, respectively. The subsequent columns present the volumes of CHE approaches: CH-COV, CH-MCD, and CH-MVE. In Fig. 3, we summarize these results by grouping the volumes calculated for all the species with each method.

**Figure 3:**
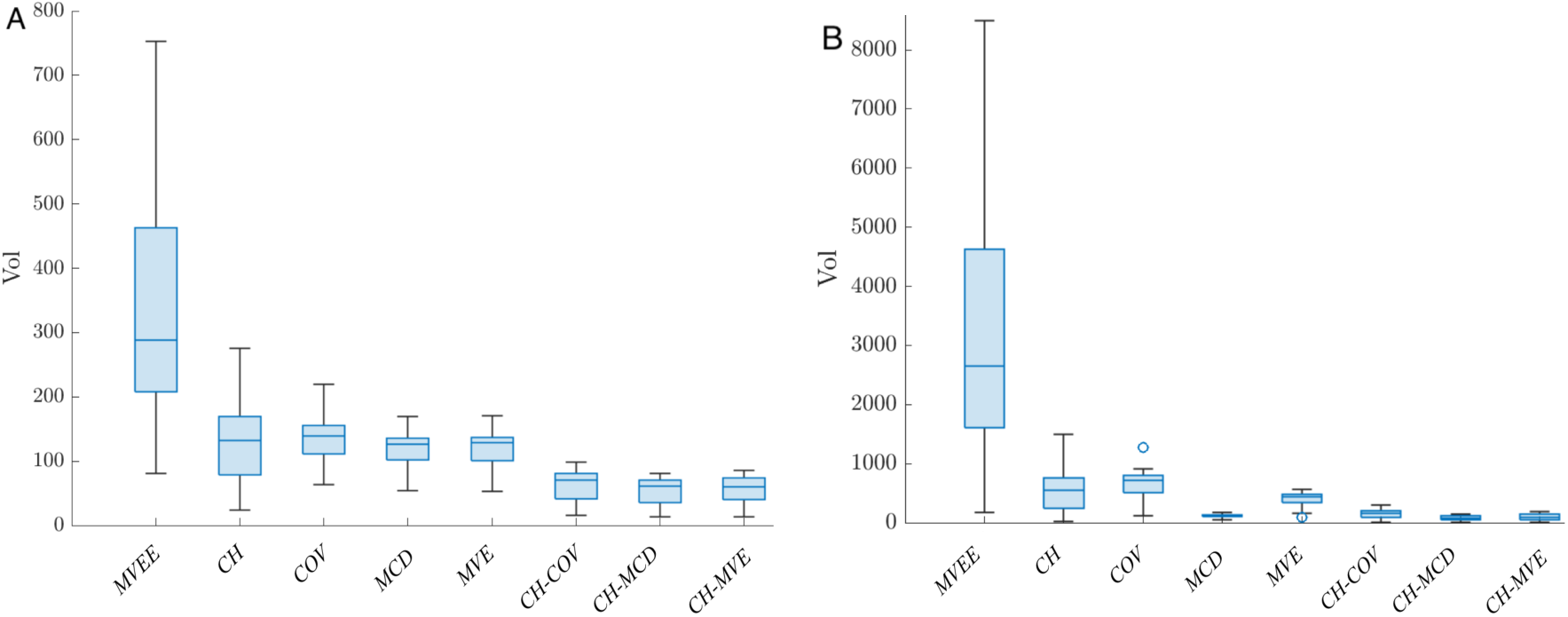
Comparison of volumes estimated for all the species and under each method for A) *n* = 3 and B) *n* = 4.

For *n* = 3, the average volumes (Vol) obtained were 339.29±193.69, 128.97±69.514, 133.7±33.05, 118.22±29.03, 62.04±24..12, 54.62±24.32 and 55.43±22.12 corresponding to MVEE, CH, COV, MCD and MVE, CH-COV, CH-MCD and CH-MVE approaches, respectively (see Fig 3-A). In the same way, average volumes were obtained for *n* = 4 with values of 3388.4±2846.4, 545.26±396.93, 667.68±233.87, 118.03±28.63, 388.82±122.42, 142.54±77.67, 79.82±42.607, 95.13±52.52 corresponding to MVEE, CH, COV, MCD and MVE, CH-COV, CH-MCD and CH-MVE approaches, respectively (see Fig. 3-B). For *n* = 3 (Fig. 3-A), the variability of the volumes estimated with CHE-approaches is smaller that the one observed for the remaining ones. For *n* = 4 (Fig. 3-B), these trends are not strictly maintained, however, the hypervolumes fitted with the CHE approach are in general smaller than the method used in the first step of the fitting process. In both cases, the MVEE approach show a higher variability and larger volumes compared to the other approaches.

### 3.2 Sensitivity analysis

In Fig. 4, we show the estimated volumes obtained with the MVEE, CH, COV, MCD, MVE and CHE approaches for different confidence levels (*p* = 0.9, 0.91, …, 0.99) and dimensions (*n* = 2, …, 7). In the left chart (A), we see that the MVEE and CH methods have fixed values (because they do not depend on *p*), so they are represented by a straight line parallel to the *x*-axis. On the other hand, the hypervolumes estimated with the COV, MCD and MVE increase in volume when the confidence level increases. In particular, the volumes calculated from enclosing envelope methods 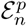 increases faster than of volumes of 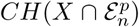 for the different approaches: COV, MCD and MVE. Furthermore, the volumes from the CHE-MVE method shows a linear trend, while the other CHE methods shows an exponential effect as the confidence level increases.

**Figure 4:**
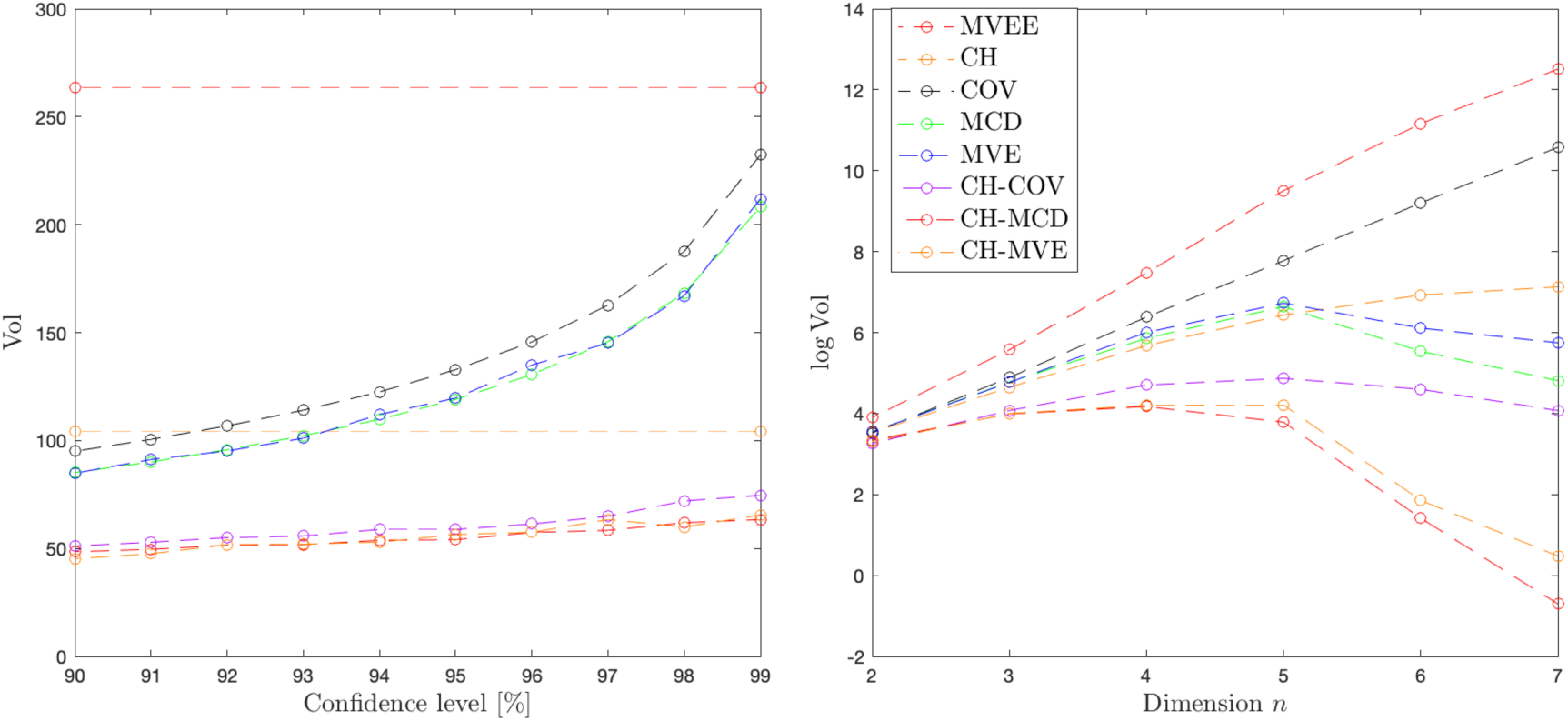
Charts showing the variation of volume with respect to A) confidence level, and B) hyperspace dimension, using the presence data of the species *B. barbastellus*.

**Figure 5:**
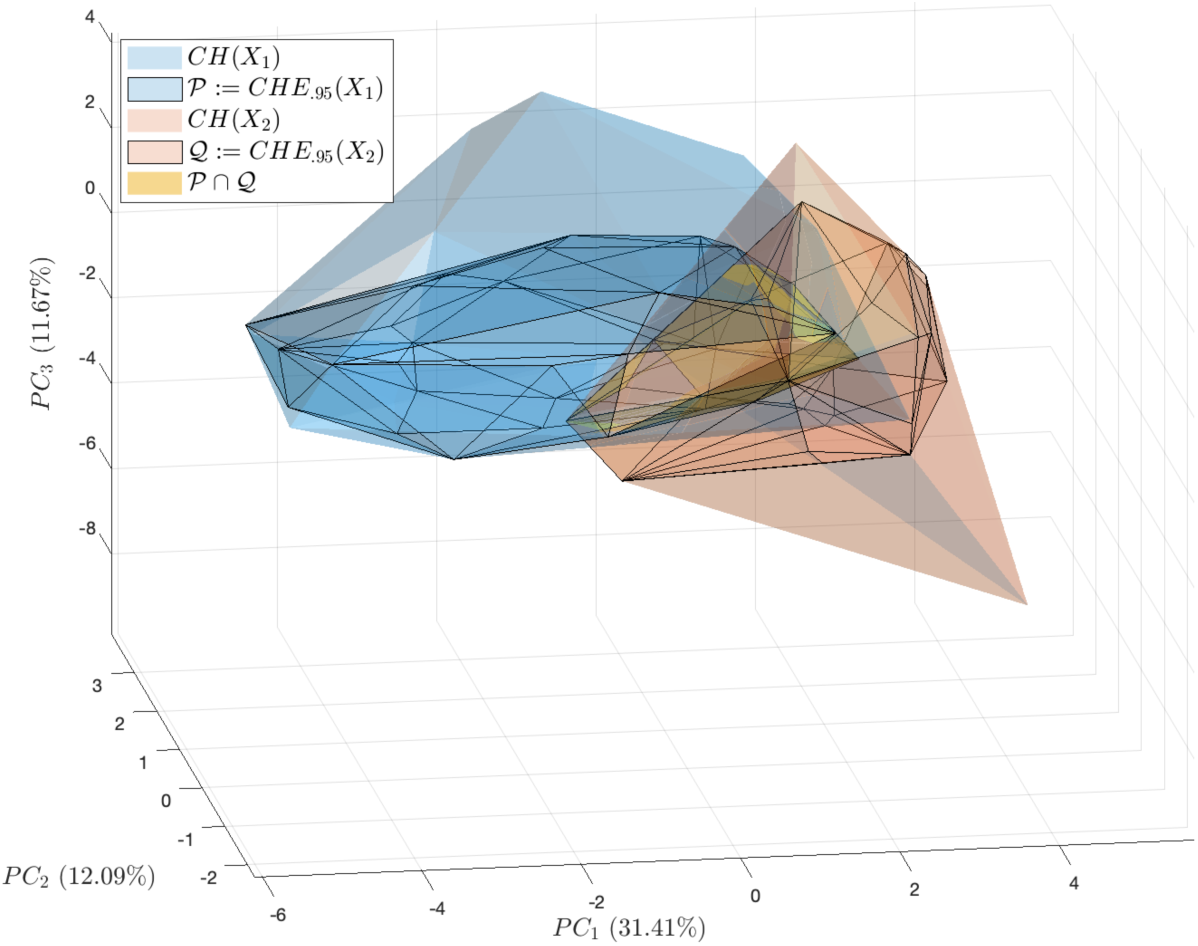
Representation of two hypervolumes 𝒫 and 𝒬, and their overlap 𝒫 ∩ 𝒬in a three-dimensional space. 𝒫 and 𝒬 were constructed with the CH-MCD method and the presence data of the species *B. barbastellus* (*X*_1_) and *E. isabellinus* (*X*_2_), respectively.

Fig. 4-B shows the effect of volume when the dimension of the hyperspace varies using a fixed value of *p* = 0.95. Because the volumes reach very large values, specially at high values of *n*„ we re-scaled the y-axis by applying the natural logarithm. We can see that the methods MVEE, COV and CH show increasing curves, while the remaining ones show a positive trend until reaching a certain dimension (*n* = 4 or *n* = 5), and then they decrease. Note also that the method CH-MVE produces the smallest volumes regardless of the value of *n*, while the method MVEE, produces the largest hypervolumes.

Finally, there is no big difference among the different methods when calculated in a space of low dimensions (*n* = 2, 3, 4), and as we include more variables, the size of the hypervolumes becomes more relevant. For example, if we wish to compare the hypervolumes of different species in a space of more than four dimensiones, we would find a higher degree of overlap among the species-specific hypervolumes when using a MVEE method and a lower degree of overlap when applying a CHE approach.

## 4 Discussion

### 4.1 Hypervolume size

We have proposed a new methodology for the estimation and construction of ecological niches that combines two of the most popular approaches used in the literature: convex hull and elliptical envelope methods, which we called the *CHE approach*. The CHE method inherits the good properties of both worlds: it fits data better (in the sense that it identifies better where the high density of observations is) like the CH does, and it is robust to outliers like the EEs. Based on our experiments, which included analyses on 25 sets of presence-only data from different bat species, the volumes estimated with our approach are in general significantly smaller than the ones obtained with baseline methods.

Theoretically, the volume of the convex hull of the set 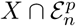 is less than or equal to the volume of the convex hull of the set *X*, because the former is included in the latter, and under the same argument is less than or equal to the volume of the set 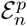, for any ellipsoid built with COV, MCD or MVE. From Fig. 3, we concluded that in general, the volumes estimated with CH-MCD approach reaches lower values compared to the other approaches, and this difference is even more pronounced when *n* = 4. Also, it is interesting to note that the MCD approach has values comparable to the CHE approach when combined with the MCD method, when *n* = 4. For both *n* = 3 and *n* = 4, the CH-MCD approach fitted hypervolumes of smaller size on average, and for *n* = 3 it leads to a lower standard deviation, which does not happen for *n* = 4, where the minimum value is reached with MCD (see Fig. 3-B).

The results obtained in Section 3.2 revealed that the volumes obtained by EE methods exceeded those obtained by the other methods (CH-COV, CH-MCD and CH-MVE) generating larger niches. This difference was more pronounced as the confidence level increased (see Fig. 4-A), except for the MVEE methods which does not depend on *p* and it. MVEE does not leave presences outside the fitted of hypervolume, in contrast to the COV, MCD and MVE. The MCD and MVE approaches are more sophisticated than the classic COV (Rousseeuw and Driessen, 1999; Van Aelst and Rousseeuw, 2009), because they were designed to protect against outliers, and produce smaller volumes (see e.g. *M. emarginatus* in Table 2, where the volume of MCD exceeds that estimated by COV). On the other hand, although the CH approach produces on average volumes smaller than the COV method (see Table 2), but similar to the ones obtained with the MCD and MVE methods, it showed a higher standard deviation, showing that outliers can dramatically increase size (and modify the shape) of the hypervolume that it generates. Furthermore, Fig. 4 shows that the volumes obtained by the methods MCD and MVE are smaller that the ones obtained with the CH method for confidence levels between approximately 90% and 92%. In contrast, the CHE approaches showed a linear trend as a function of the confidence level and overall produced the smallest hypervolumes.

We also showed that the methods MVEE, COV and CH tend to produce larger hypervolumes as *n* increases (although a stabilization of CH is shown from dimension *n* = 5; see Fig. 4-B). Given that the hyperspace was defined with principal components, as *n* increases, the variability in that dimension decreases (see Fig. S1), and therefore these approaches are not capturing this decrease in an adequate way. The MCD, MVE and CHE methods showed increasing volumes for *n<* 5 and an inflexion point at *n* = 5, where the volumes started to decrease. This change was more prominent for the CH-MCD and CH-MVE methods, showing that they better capture the variability of the principal components.

### 4.2 Hypervolume overlap

The intersection or the overlap region of two *n*-dimensional hipervolumes plays an important role in ecology in a number of applications (Cavalcante et al., 2020; Donoghue and Edwards, 2014; Guisan et al., 2014; Peterson et al., 2013; Rissler and Apodaca, 2007). In general, there are not known formulas for the intersection of *n*-dimensional ellipsoids, and therefore, approximation methods that involve great computational effort are commonly used (Qiao et al., 2016; Rabiei and Saleeby, 2021; Swanson et al., 2015). In this context, we emphasized already that the CHE approach produces polyhedral hypervolumes that can be represented in the form of a set of inequalities. This is a central feature because it makes easier to calculate the intersection between two hypervolumes. If 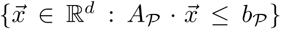 and 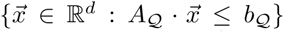 are the polyhedral representations of the hypervolumes 𝒫 and 𝒬 respectively, then 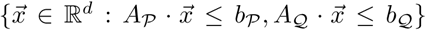 is the polyhedral representation of the hypervolume overlap: 𝒫 ∩ 𝒬, which is efficiently quantified with our methodology (see *intersect function* in Multi-Parametric Toolbox (Herceg et al., 2013)).

The polyhedra whose faces are delimited by black lines represent the hypervolumes built with the CHE approach (CH-MCD) for *p* = 0.95, and using the presences of the species *B. barbastellus* (blue polyhedra) and *E. isabellinus* (red polyhedra), while the shaded (larger) polyhedra were fitted with the classic CH approach for the same two species. In yellow, we highlighted the overlap of both of the hypervolumes fitted with the CHE approach. The niche overlap between the hypervolumes fitted with the CHE method is smaller (Vol(𝒫 ∩ 𝒬) = 7.42) than the overlap between the convex hulls (Vol(*CH*(*X*_1_) ∩ *CH*(*X*_2_)) = 22.92). Therefore, when using the CH method to approximate the realized niche, there is an overestimation of niche overlap and the intersection of the niches probably contains irrelevant regions of the environmental space. The CH-MCD method excludes environmental conditions that are poorly represented in the sample of presence points, giving a more meaningful representation of niche overlap.

When we project these hypervolumes to geography as models for the realized niche, there is also a decrease in the number of cells that are predicted to be part of the realized niche of each species (see Figures S2 and S3 in the Supplementary Material) when comparing the CH against the CHE approach. As a consequence, the area predicted as suitable for both species changes depending on the method used to estimate the niches. In Fig. 6-A, we see that when we use the CH approach to fit the realized niches of *B. barbastellus* and *E. isabellinus*, the overlap among the fitted hypervolumes is represented by a larger number of yellow cells (2,255 cells), compared to the yellow area in Fig. 6-B (1,158 cells) obtained from the overlap of the CHE. Broadly, both the CH and CHE approaches always produce overlaps that contain at least one point of the observations, an aspect that is not ensured with niches built with EE methods. However, since the hypervolumes generated by the CHE approach shrink towards the centroid of the ecological niche (when we decrease the confidence level), they are more likely to contain a higher number of observations of the species and, therefore, they may be a better representation of the realized niche of the species.

**Figure 6:**
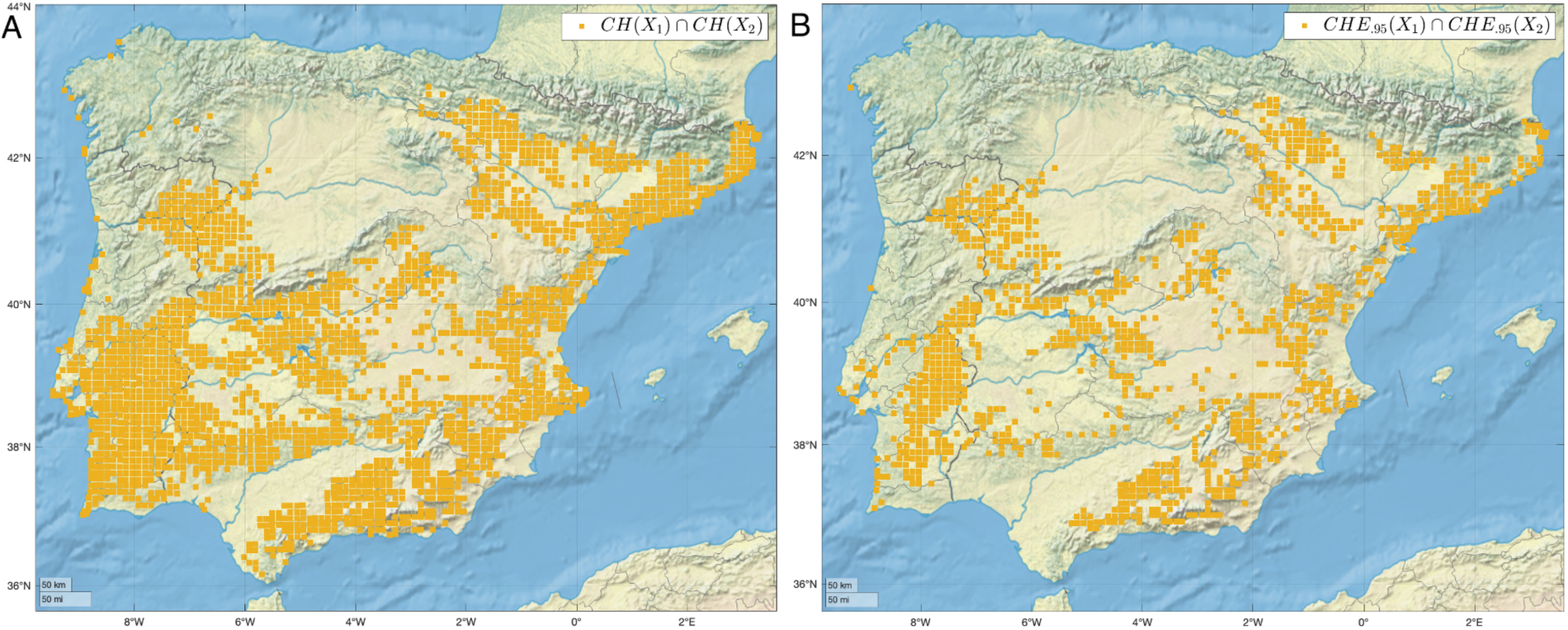
Niche overlap for the species *B. barbastellus* and *E. isabellinus* mapped into the study area. A) The yellow cells correspond to the overlap between the hypervolumes fitted with the CH approach. B) The yellow cells represent the overlap between the hypervolumes fitted with the CHE approach (CH-MCD).

## 5 Conclusions

The CHE procedure allows the construction of niche-hypervolumes that can be used as models for the realized niche of a species, as shown for the selected Iberian bat species, by combining the convex hull and elliptical envelope methods. This approach fits polyhedral hypervolumes based on presence-only data, and allows to define regions of iso-suitability, as a function of significance statistics inherited from the Mahalanobis distance model or from the robust estimators obtained with methods such as MCD and MVE.

Our results showed that our niche modeling approach better suits the presences of the species, is versatile, and preserves the interpretability of classical convex and elliptical envelope approaches, and remaining aligned with the definition of Hutchinson’s niche (Hutchinson, 1957).

The ability to calculate niche overlap has received much attention in the last decade, and is a desirable property of niche modeling approaches, the estimation of which is directly included in the Sorensen-Dice and Jaccard similarity metrics (Mammola, 2019; Cerdeira et al., 2018). We were able to corroborate that convex hulls hypervolumes are sensitive to outliers, and might overestimate the realized niche of a species as well as the intersection of the niches of two or more species, therefore producing incorrect similarity measures. The CHE approach can produce more precise and more robust estimates of both hypervolumes that represent the species realized niche and niche overlaps, although more research in that direction is necessary.

