## Supplementary Material for "A new method to estimate the ecological niche through *n*-dimensional hypervolumes that combines convex hulls and elliptical envelopes"

### Suplementary Material

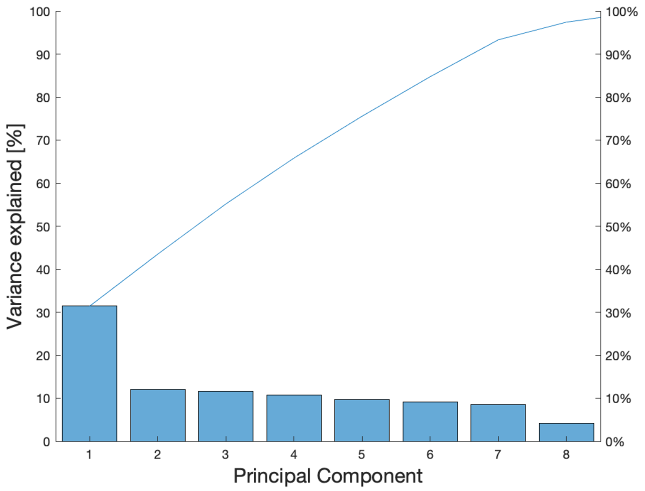


Figure S1: Percentage of variability explained by each principal component calculated with the variables listed in Table 1.

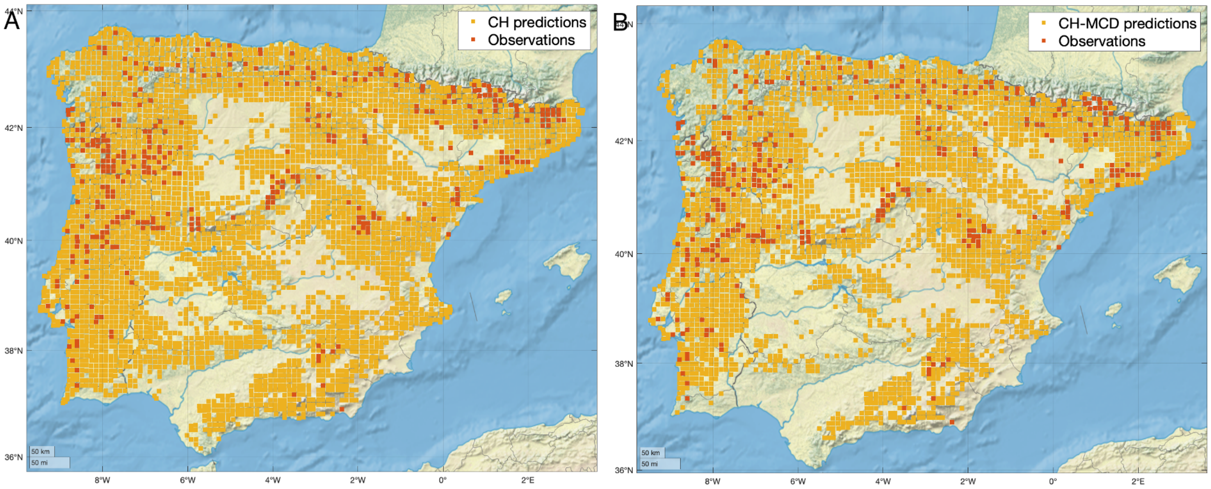


Figure S2: Projection of hypervolumes to geography for the species *B. barbastellus.*

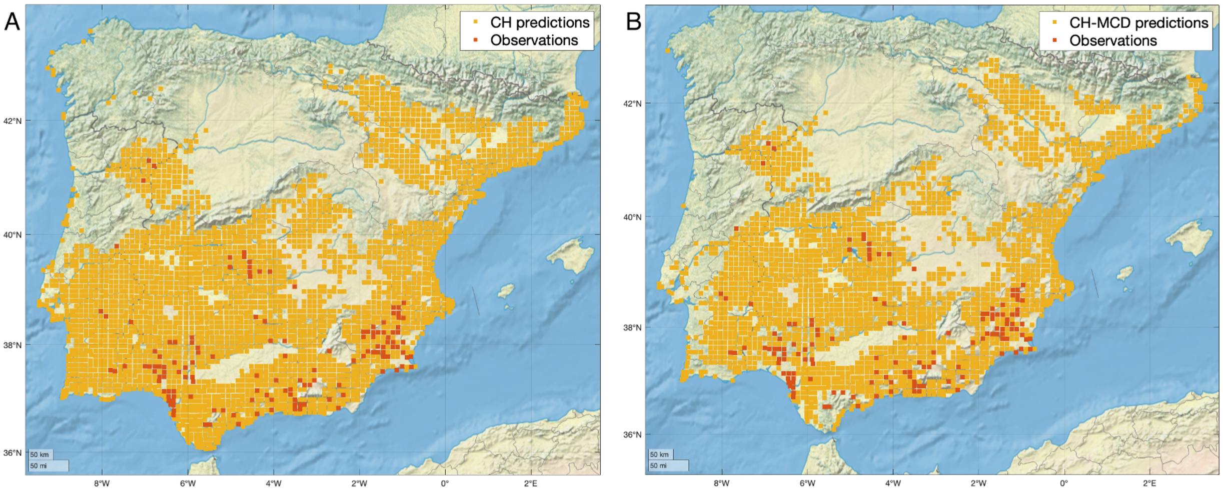


Figure S3: Projection of hypervolumes to geography for the species *E. isabellinus*.
